# ALM enables contextual decision-making via dynamic reconfiguration of local circuits

**DOI:** 10.1101/2025.09.11.675319

**Authors:** Jia Shen, Nuttida Rungratsameetaweemana, Prayshita Sharma, Darcy S. Peterka, Herbert Zheng Wu, Michael N. Shadlen

## Abstract

Cognitive operations often require flexible implementation of stimulus–response contingencies, depending on context. We developed an olfactory task in which mice learned to associate a test odor with a directional lick response, conditional on a preceding context odor drawn from a different odor set. Two-photon imaging revealed that anterior lateral motor cortex (ALM) contains distinct populations encoding context, test odors, and choice. Optogenetic silencing during the context and delay periods impaired performance, suggesting that ALM contributes to configuring the appropriate contingency. Although context odors that instructed the same mapping were represented by separate populations, their influence converged at the level of choice-selective neurons. A subpopulation of these neurons exhibited dual selectivity for context and choice, forming what we term “contingency neurons.” These findings suggest that ALM supports flexible behavior not by abstracting over context cues, but by dynamically reconfiguring local circuits to route sensory input to the appropriate motor output.

## Introduction

The ability to flexibly integrate sensory, contextual, and internal information to guide behavior is a core function of cognitive control, enabling organisms to override fixed stimulus–response mappings^1,2^ (SR-mappings from here on). Such flexibility permits the same stimulus to elicit different actions depending on task demands or environmental conditions. For example, a driver in the United States enters a car from the left, whereas in the United Kingdom, entry occurs from the right. This context-dependent modulation of behavior extends the repertoire of possible actions and is essential for adaptive decision-making. Notably, deficits in behavioral flexibility are observed across several neuropsychiatric conditions, including autism spectrum disorder, attention-deficit/hyperactivity disorder, and dementias, where actions tend to become rigid or resistant to change^3,4^. Although the behavioral phenomena are well described, the underlying neural circuit mechanisms that support flexible SR-mapping remain incompletely understood.

Emerging evidence implicates the secondary motor cortex (M2), particularly its anterior-lateral region (ALM), in establishing flexible sensorimotor associations^5,6^. ALM neurons have been shown to maintain decision-related information—such as an intention to lick left or right—through persistent activity during delay periods in cued response tasks^7–10^. In a recent study, Wu et al. ^6^ employed an olfactory delayed match-to-sample (DMS) task to investigate context-dependent decision-making. They discovered odor-selective neurons in superficial ALM that supported a trace of the sample odor and neurons in deeper layers that supported the directed-lick report (left or right for match or non-match, respectively). Optogenetic inhibition of ALM during the sample and delay periods impaired behavioral performance, thus supporting a causal role of ALM in the DMS task.

Partial inactivation experiments, however, led Wu et al to propose that the necessary role of ALM was not to maintain a trace of the sample odor for subsequent comparison to the test odor, but rather to establish the appropriate SR-mapping between the test odor (stimulus) and directed lick (response). They reasoned that such a mechanism (e.g., dendritic gating) could prepare the SR-mapping in the brief window of normal activity furnished by the partial inactivation protocol. However, so long as the same set of odors are presented as sample and test odors, it is difficult to rule out the possibility that a low-level comparison mechanism, such as familiarity detection^11,12^, plays a role.

To address this limitation, we developed a variant of the DMS task using four distinct odorants. One of two odors, limonene and methyl butyrate (hereafter odors C and D) were presented first, like the sample odors in DMS. After a delay period, one of a different set of two odors, pinene and hexene (odors A and B) were presented second, like the test odors in DMS. In this “CDAB” task, the first and second odors are never matched, eliminating the confound of same-different comparisons. Accordingly, we refer to the first odor as the context odor and the second odor as the test odor. Using this paradigm, we identified (1) a population of layer 2/3 ALM neurons that exhibit selective responses to contextual odors, and (2) a requirement for ALM activity during the context and delay periods for accurate task performance. By combining the original DMS and the new CDAB tasks, we uncovered a population of ALM neurons that encode context-dependent contingencies. Although odors associated with the same SR-mappings (e.g., A & C or B & D) were represented by largely non-overlapping sensory populations, they appeared to converge on a shared set of choice-selective neurons. These findings suggest that ALM integrates distinct contextual inputs to dynamically configure local circuitry, thereby enabling flexible, context-dependent action selection.

## Results

We used three task variants: (*i*) the original delayed match-to-sample (DMS) task, referred to as the ABAB task; (*ii*) a novel task with distinct context odors, referred to as the CDAB task; and (*iii*) an eight-trial-type (8T) task, which randomly interleaves trials from variants *i* and *ii*. In all tasks, head-fixed mice learned to associate a test odor (A²ⁿᵈ or B²ⁿᵈ) with a left or right lick response, depending on a preceding context odor (X¹^ˢᵗ^) ^1^shorth-hand for generic 1^st^ odor) notations for), separated by a delay (**Fig. 1a**). In the ABAB task, both context and test odors were drawn from the same set {A, B}, as in Wu et al.^6^. In the CDAB task, context odors C¹^ˢᵗ^ and D¹^ˢᵗ^ instructed the same SR-mapping as A¹^ˢᵗ^ and B¹^ˢᵗ^, respectively (**Fig. 1b**).

**Figure 1.**
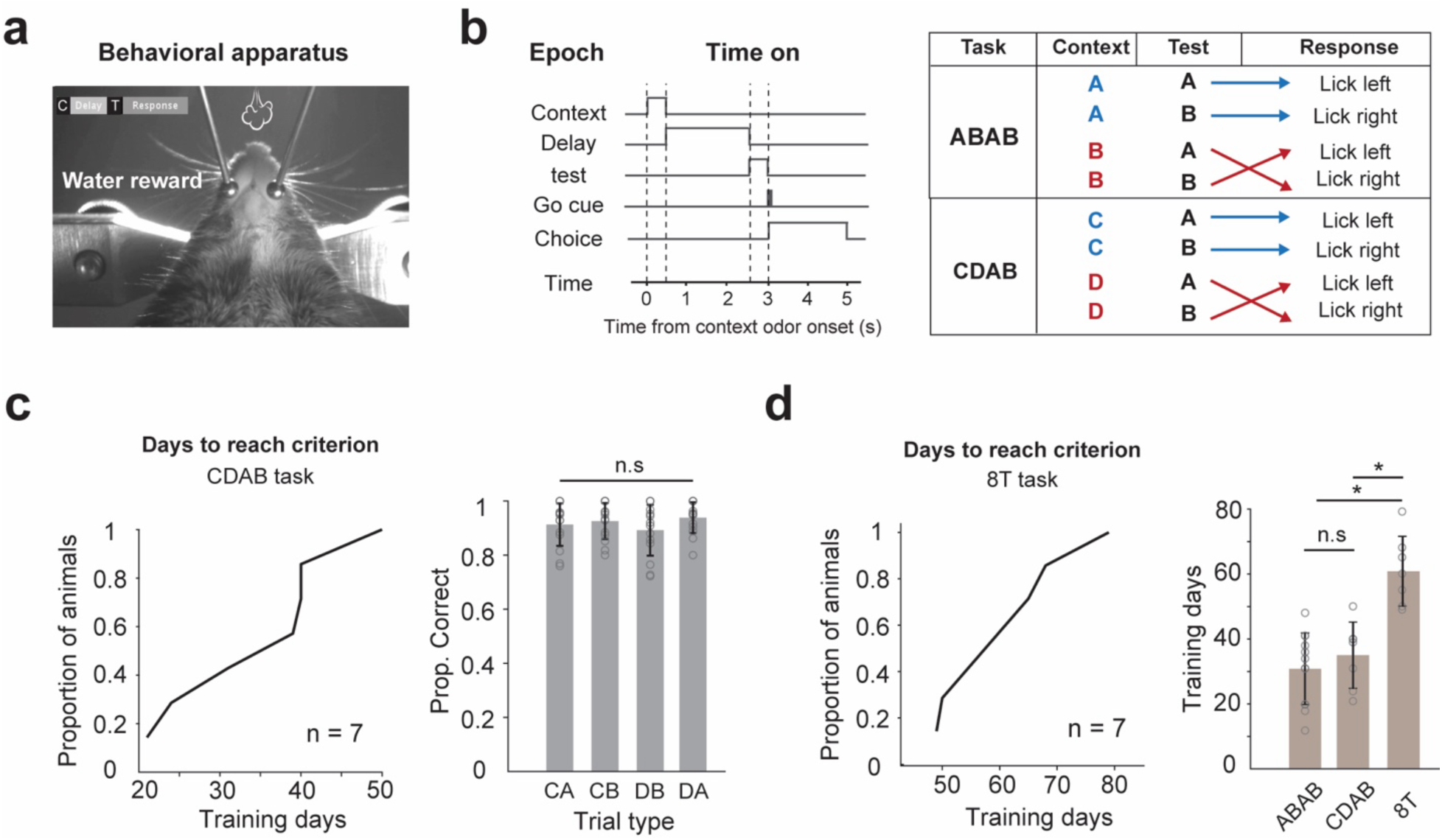
Task structure and behavioral performance. **a.** Behavioral apparatus. Photo contains one representative video frame taken during the context epoch. The camera is below the mouth, pointing upward. **b.** Task structure. *Left*, two odors, separated by a delay, are presented on each trial. The first signals the context; the second is the test stimulus. *Right*, the association between test stimulus and rewarded lick response depends on the context. The ABAB task is also known as *delayed match to sample*. The CDAB task is structured identically to ABAB, substituting context odor C for sample odor A and context odor D for sample odor B. Note that all CDAB trials are non-match. The second odor in both tasks is the test odor A or B. **c.** Behavioral performance in the CDAB task. *Left*, training days required to achieve criterion performance on the CDAB task (n = 7 mice, 35 ± 4 days). *Right*, performance of each trial type in the CDAB task (n = 14 sessions; n.s., not significant, repeated-measures ANOVA). **d.** Behavioral performance on the 8T task. *Left*, days required to achieve criterion performance (n = 7, 60 ± 4 days). *Right*, mean training days in the DMS, CDAB, and 8T tasks (\**p* < 0.05, Wilcoxon signed-rank test).

In the CDAB task, the association between the rewarded port and the test odors depended on a context odor. After context odor C¹^ˢᵗ^, odor A²ⁿᵈ required a left lick and B²ⁿᵈ required a right lick; under context D¹^ˢᵗ^, the rewarded associations were reversed (**Fig. 1b**). Context and test odors were separated by a two-second delay. Mice reached performance criterion in the CDAB task after 35 ± 4 days of training, comparable to the ABAB task (**Fig. 1c–d**; **Supplementary** Fig. 1b). Performance accuracy was consistent across trial types, indicating that animals generalized across task conditions (**Fig. 1c**; **Supplementary** Fig. 1a). Mice trained on the interleaved 8T task required significantly longer to reach criterion (61 ± 4 days; Fig. 1d), consistent with the increased task complexity. We next describe the neural substrates that support this process, comparing ALM activity during CDAB performance to that previously reported in the ABAB task.

### ALM encodes context cues, test odors, and choice during the CDAB task

To identify task-related neural representations, we performed two-photon Ca²⁺ imaging of pyramidal neurons in ALM during the CDAB task (**Fig. 2a**). The responsive neurons comprise three functional types that encoded contextual information, test odors, and choice, respectively (**Fig. 2b–d**). Approximately 15% ± 2% of neurons responded selectively to the context odors C¹^ˢᵗ^ or D¹^ˢᵗ^ (**Fig. 2f**). These neurons were primarily located in superficial layer 2/3, similar to those observed in the ABAB task, and displayed heterogeneous latencies and response dynamics (**Fig. 2e–g**; **Supplementary** Fig. 2a). As a population, they maintained a persistent representation of the context throughout the delay^9,15^ (**Fig. 2e**). A smaller proportion (8.5% ± 1.5%) responded selectively to the test odor (A²ⁿᵈ or B²ⁿᵈ) regardless of the context (**Fig. 2c**; **Supplementary** Fig. 2). The largest group (40% ± 8%) was selective for the choice—left or right licking. They were largely unresponsive during the context and delay periods (**Fig. 2d–e**; **Supplementary** Fig. 2a). The distribution and prevalence of these three neuron types in the CDAB task closely matched those previously reported in the ABAB task (**Fig. 2g**), suggesting that ALM may implement similar representational motifs across different task contexts.

**Figure 2.**
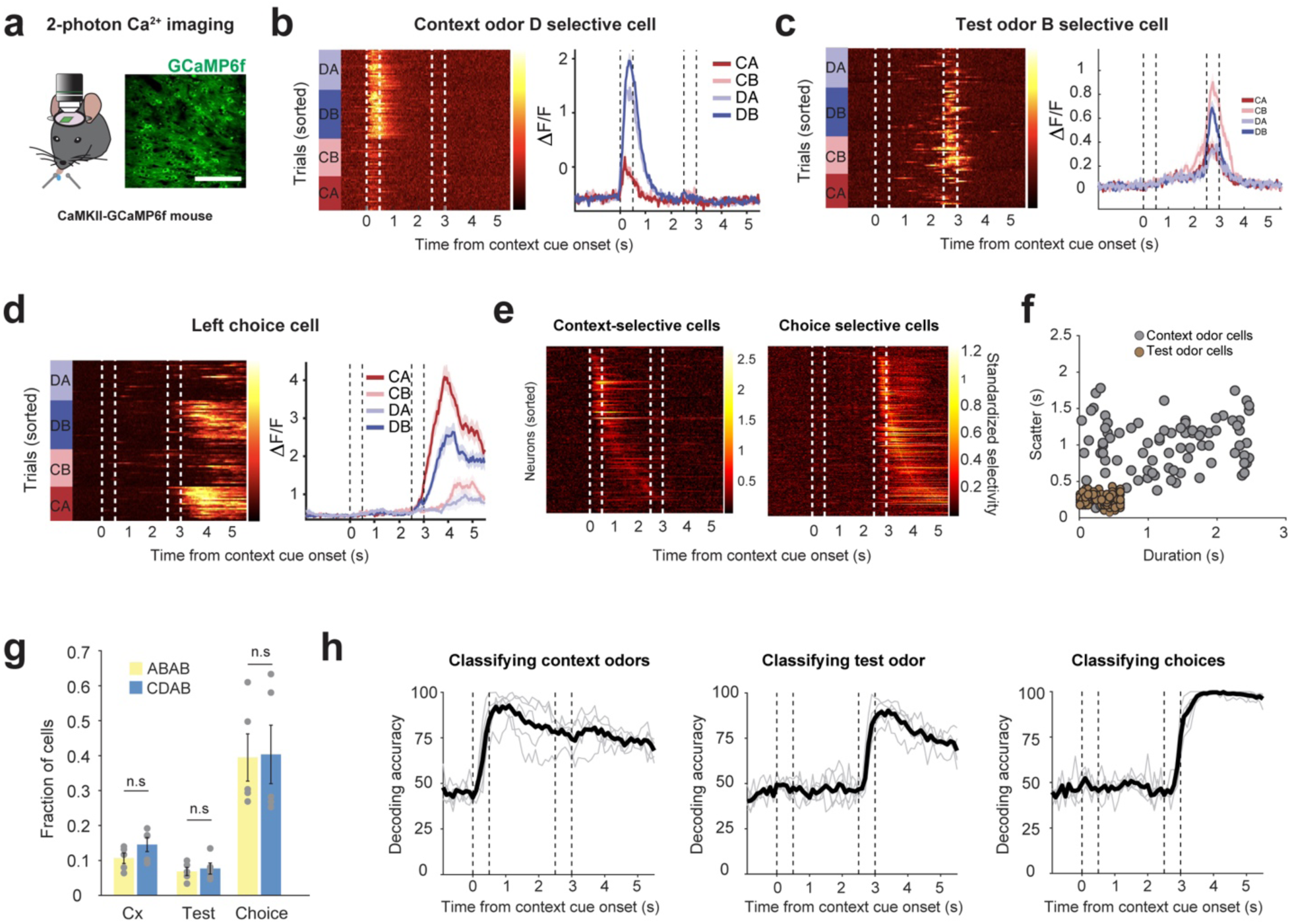
ALM neurons encode context, test, and choice in the CDAB task. **a.** 2p Ca^2+^ imaging in a GCaMP6f mouse line. The micrograph is a representative FOV from the 2p imaging. Scale bar: 50 µm. **b–d.** Representative examples of the three cell types. *Left display* in each panel is a heatmap of Ca^2+^ responses from all trial types; *right display* shows averaged Ca^2+^ responses grouped by trial type. (*b*) Context Odor-D selective cell; (*c*) Test odor-B selective cell; (*d*) Lick-left choice selective cell. **e**. *Left*, all context selective cells sorted by the time of their peak standardized selectivity; *Right*, all choice selective cells (same sorting schema). **f.** Scatter plot. Each symbol shows the average duration of the neuron’s response (abscissa) and the variability in the timing of the response during the context/delay period (ordinate; the scatter metric is defined in Methods). Unlike the stereotyped dynamics of test odor-selective neurons (brown), context odor-selective neurons are heterogeneous. **g.** Distribution of functional cell types is similar for mice trained to perform the ABAB task (data from Wu et al) the CDAB task (present study; n = 5 mice; n.s., not significant, Wilcoxon signed rank test). **h.** The performance (held-out trials) of an SVM classifier trained to identify the context from the population calcium activity. Thin traces are individual sessions; thick line is the mean.

### ALM activity during context and delay is required for task performance

Previous work showed that suppressing ALM activity during the context and delay periods impaired performance on the ABAB task^6^. Based on the similar task structure and behavioral demands, we hypothesized that ALM also plays a causal role in the CDAB task. To test this, we trained VGAT-ChR2 mice on the CDAB task and optogenetically activated inhibitory neurons bilaterally in ALM during the context and delay epochs. The laser ramped off at the end of the delay to avoid interfering with licking execution (**Fig. 3a**; see Methods). Inhibiting ALM during the context and delay periods caused pronounced performance deficits (**Fig. 3b**), whereas partial inactivation, limited to the context odor and early delay, produced intermediate deficits (**Fig. 3c**). Suppression during the test and choice epochs abolished licking behavior altogether (data not shown), consistent with ALM’s established role in motor execution^7,9,16,17^. These results support the hypothesis that intact ALM activity during the context and delay periods is necessary to configure the circuit appropriately so that the test odor elicits the correct response.

**Figure 3.**
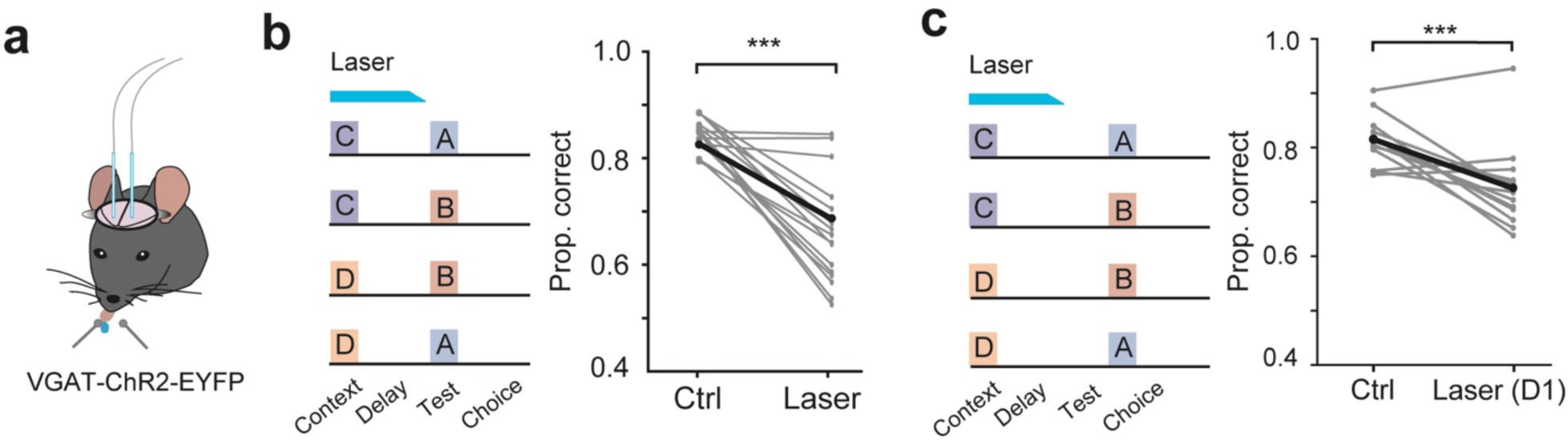
Inactivation of ALM impairs performance on the CDAB task. **a.** Optical inhibition of ALM in both hemispheres through excitation of GABAergic neurons (same mouse line as in Wu et al and identical protocol). The laser is on through the context epoch and most of the delay. It is ramped off over the last 250 ms preceding the test odor. **b.** The choice accuracy was impaired in the laser-on trials. than control trials (n = 35 sessions, \*\*\**p* < 0.001, Wilcoxon signed-rank test, comparison to control trials). **c.** Effect of inactivation during the context and first 1.5 s of the delay epoch (n = 14 sessions, \*\*\**p* < 0.001, Wilcoxon signed-rank test). Data from each session is shown in gray; black symbols are the means over sessions.

### Context odors are represented by distinct ALM populations

To investigate how this context-dependent configuration might be implemented, we next examined neural activity in the 8T task, which interleaves ABAB and CDAB trial types. This design enables us to dissociate odor identity from the context those odors convey. Specifically, odors A¹^ˢᵗ^ and C¹^ˢᵗ^ both instruct the same SR-mapping (Context 1, or Cx1: A²ⁿᵈ → lick left; B²ⁿᵈ → lick right), while B¹^ˢᵗ^ and D¹^ˢᵗ^ signal the alternative mapping (Cx2: A²ⁿᵈ → lick right; B²ⁿᵈ → lick left). Most X¹^ˢᵗ^ odor–selective neurons responded to only one of the four context odors, but a small subset responded to two. If ALM generalized across odors that signal the same context—for example, A¹^ˢᵗ^ and C¹^ˢᵗ^ (Cx1)—we would expect this overlap to preferentially group same-context odor pairs^18^. To test this, we asked whether neurons responsive to multiple X¹^ˢᵗ^ odors tended to be selective for odors associated with the same SR-mapping.

We found little evidence for such grouping. Although some neurons responded to both A¹^ˢᵗ^ and C¹^ˢᵗ^, or to both B¹^ˢᵗ^ and D¹^ˢᵗ^ (**Fig. 4a–e and Supplementary** Fig. 3a **&. b)**, a similar number responded to odor pairs associated with different mappings (e.g., A¹^ˢᵗ^ and D¹^ˢᵗ^; **Fig. 4a–c, f, and g**; **Supplementary** Fig. 3c **& d**). Example neurons illustrated in **Fig. 4d–e** show selective responses that align with both same-context and opposing-context pairings. Quantitatively, the overlap between same-context odor pairs was not significantly greater than the overlap between different-context pairs (**Fig. 4c**; permutation test). These results suggest that X¹^ˢᵗ^ selective neurons in ALM do not generalize across odors that signal the same behavioral context. This conclusion is further supported by the following decoding exercise.

**Figure 4.**
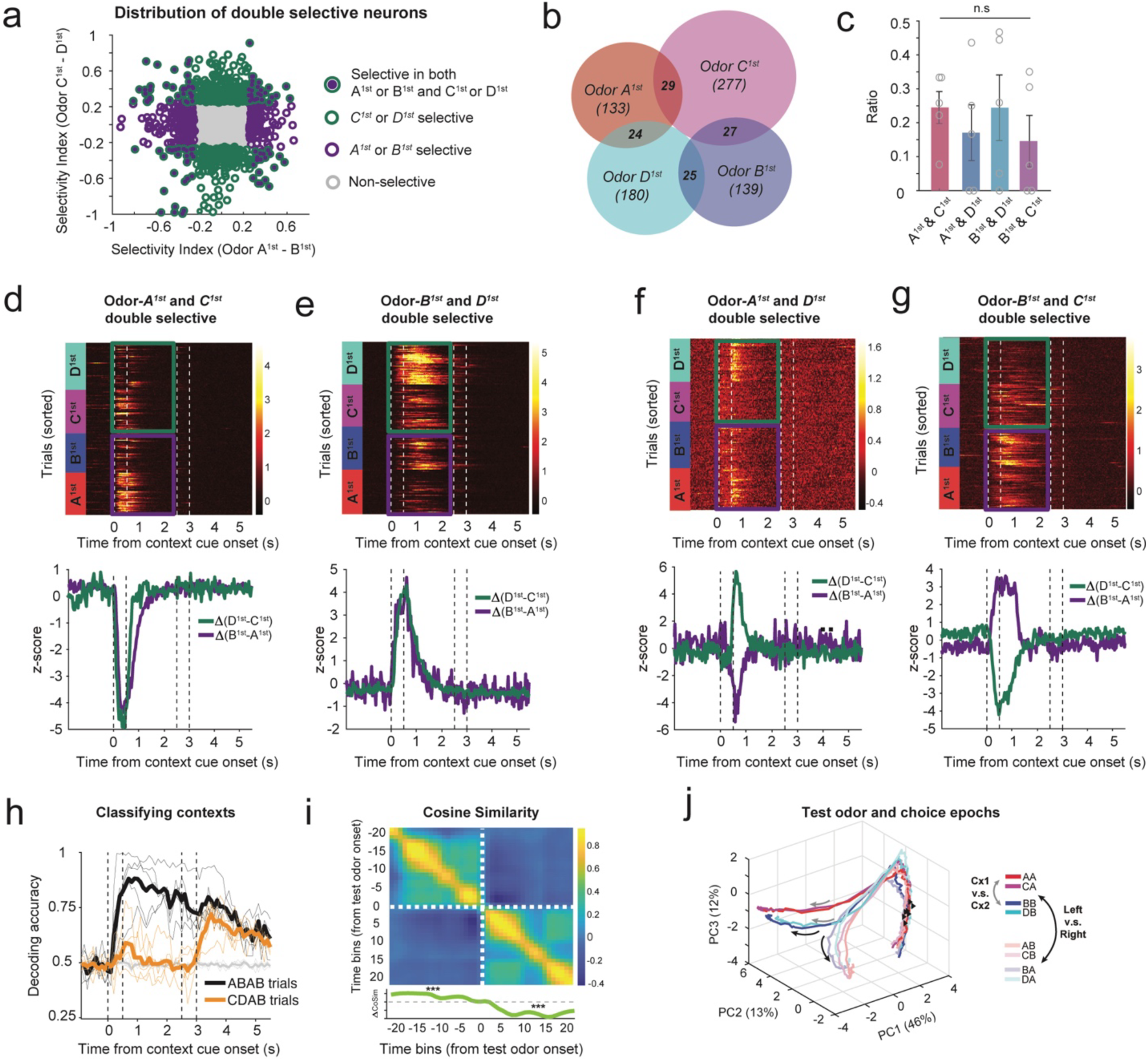
Context odors are represented by distinct ALM populations. **a.** Scatter plot shows each neuron’s selectivity index (SI) for context odors C vs. D (CDAB trials; ordinate), plotted against its SI for A^1st^ vs. B^1st^ (ABAB trials; abscissa). **b.** Venn diagram shows the overlap of neurons with statistically significant responses to each of the four X^1st^ odors. Doubly selective neurons are tabulated at the intersections. **c.** The number of doubly selective neurons that signal the same context is similar to the number that signal different contexts (n = 5, n.s. not significant, permutation test). **d–g.** Examples of doubly selective neurons. Upper displays are heat maps of the activity in all trials; rows are single trials. Green and purple rectangles outline CDAB and ABAB trials, respectively. Lower displays are averaged standardized activity in the heatmap above. Panels **d** & **e** show examples of doubly selective neurons for first odors that signal the same context. Panels **f** & **g** show examples of neurons doubly selective for first odors that signal different contexts. **h.** Accuracy of an SVM classifier, trained to identify context from population activity (similar to Fig. 2h). Here the decoder is trained on ABAB trials. Training is conducted at successive time bins (100 ms). Accuracy is best when testing is restricted to held out data from ABAB trials (black trace). The classifier fails to exceed chance performance on CDAB trials (orange) in the context and delay epochs. As in Fig. 2h, Decoding of context in the choice epoch exceeds chance, and here classification accuracy of CDAB trials is similar to ABAB trials. **i.** Heatmap depicts Cosine-similarity of pairs of SVM-derived weight vectors at the times indicated along the ordinate and abscissa, respectively. CoSim vals in quadrants 1 and 4 correspond to the similarity between weight vectors derived in the choice epoch; CoSim vals in quadrants 2 and 3 correspond to the similarity between weight vectors derived in the context/delay epoch (see Methods). The origin marks the time of the test odor. *Bottom panel*, for t<0, ΔCoSim is the difference between quadrant 2 column averages and quadrant 3 column averages. For t>0, ΔCoSim is the difference between quadrant 1 column averages and quadrant 4 column averages (\*\*\**p* < 0.001, permutation test, see Supplementary Fig. 3f and Methods). **j.** Neural trajectory depicted in a 3D state space defined by the first three principal components, which explain 71% of the variance in the calcium signals across time-aligned trials. Note the divergence of traces based on choice (black arrows) and a second bifurcation reflecting context (gray arrows).

We trained a support vector machine (SVM) classifier to decode the identity of the context odor using population activity during the ABAB trials. The model accurately distinguished A¹^ˢᵗ^ from B¹^ˢᵗ^ on held-out ABAB trials, but failed to decode C¹^ˢᵗ^ and D¹^ˢᵗ^ on CDAB trials using activity from the same epochs (**Fig. 4h**). This cross-generalization failure reinforces the conclusion that same-context odors are represented by distinct neural populations. We next show that the representation of context occurs in the activity of neurons that organize the directed-lick response to the test odor.

### Generalization of context at the level of choice neurons

Figure 4h also shows that SVM classifiers, trained and tested after onset of the test odor (i.e., during the choice epoch), accurately decoded context—even though most X¹^ˢᵗ^-selective neurons were inactive during this time. This capacity did not depend on which trial set was used for training. A classifier trained on CDAB trials also decoded the context on ABAB trials (**Supplementary** Fig. 3e). Examination of the SVM weight vectors across time in the trial revealed an abrupt shift after the test odor was presented (Fig. 4i). During the context and delay periods, the classifier relied on a stable set of neurons, but after test odor onset, a new set of weights emerged, indicating that different neurons supported decoding in the choice epoch (**Supplementary** Fig. 3f). Importantly, in the choice epoch, the model trained on A¹^ˢᵗ^ vs. B¹^ˢᵗ^ could decode C¹^ˢᵗ^ vs. D¹^ˢᵗ^ with comparable accuracy—despite the lack of generalization earlier in the trial.

To visualize how context information is represented at the population level, we performed principal component analysis (PCA) on ALM activity. Using Ca^2+^ response data from the context and delay epochs, we found that the first three principal components (PCs) captured 72% of the variance across trials (**Supplementary** Fig. 3f). However, the trajectories for odors that signaled the same context—for example, A¹^ˢᵗ^ and C¹^ˢᵗ^—remained largely distinct in this low-dimensional space, consistent with the idea that ALM maintains separate representations for different context odors.

We then performed PCA on neural activity during the test and choice epochs. In this later phase, trajectories diverged sharply based on the animal’s motor response (Fig. 4j). Strikingly, trials associated with the same choice but different contexts further diverged based on the context identity (**Supplementary** Fig. 3h). In other words, within each motor response category (left or right lick), trajectories branched according to the preceding context odor, suggesting the population activity during the choice epoch retains the representation of context from earlier in the trial. These findings support the interpretation that convergence of odor representations into a categorical representation of context occurs during the choice epoch and is not evident earlier in the trial. They also corroborate the classification results, which revealed a shift in the neurons carrying information about context at this time.

### Some choice-selective neurons encode mapping contingencies

Task-related activity during the choice epoch was dominated by neurons selective for lick direction, responding preferentially for left or right choices, regardless of the preceding context or odor sequence (Fig. 5a**–b**). However, the decoding and the PCA results suggested that context information remains accessible during this epoch, raising the possibility that some choice-selective neurons might also reflect contextual input. Indeed, in the 8T task, we identified a subpopulation of ALM neurons with dual selectivity. They were selective for both both the choice direction and the context in which that choice was made. These “contingency neurons” were activated only under specific context–choice combinations, effectively encoding the SR-mapping itself (Fig. 5c**–g**).

**Figure 5.**
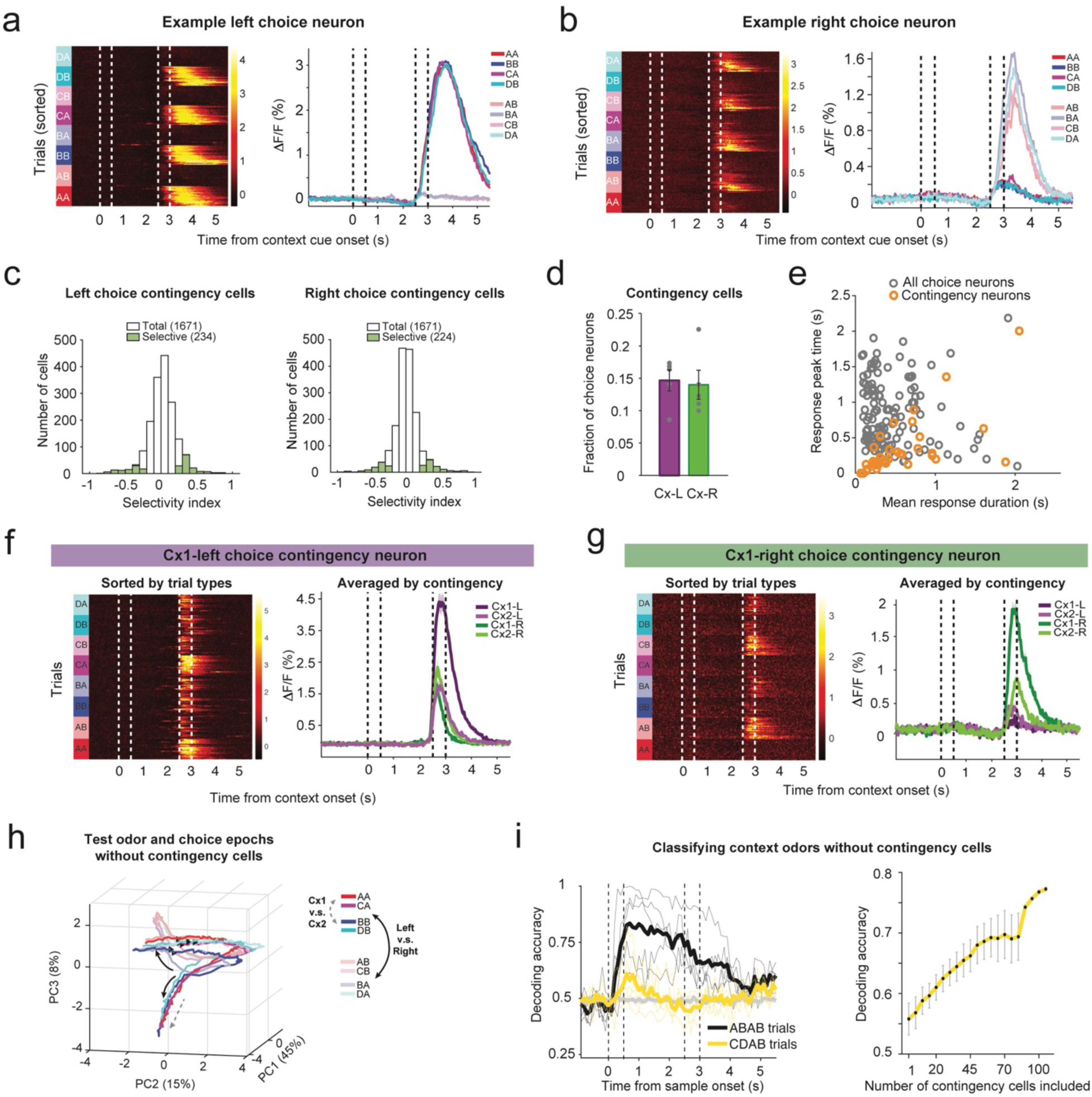
Contingency neurons link context and choice to implement flexible SR-mappings. a & b. Left displays are heatmaps showing single trial Ca^2+^ responses. Right displays are trial averaged Ca^2+^activity over trials, as a function of time. Data are from the 8T task. **a.** Representative left-choice neuron. **B.** Representative right-choice neuron. Trials are grouped by trial type (color); saturated colors indicate the rewarded directed-lick is left. **c.** Histogram highlighting the small fraction of directed-lick neurons that are also selective for context, termed “contingency neurons”. **d**. Breakdown by lick-direction preference. **e**. Contingency neurons respond earlier and more transiently than other directed-lick neurons. Peak time and duration of Ca^2+^ responses of contingency neurons (orange) and all choice neurons (grey). Response peak time: 0.81 ± 0.04 s for all choice neurons and 0.32 ± 0.02 s for contingency cells (\*\*\**p* < 0.0001, Wilcoxon signed rank test). Response duration: 0.38 ± 0.02 s for all choice neurons and 0.44 ± 0.03 s for contingency cells (n.s. *p* > 0.05, Wilcoxon signed rank test). **f.** Representative Cx1-left choice neuron. Left, Ca^2+^ responses in all trials, sorted by trial type. Right, sorted by SR-mapping (Cx1-L, Cx2-L, Cx1-R, Cx2-R). **g.** Example Cx1 right choice neuron. **h.** Neural trajectory in a 3D state space during test odor and choice epochs without the contingency neurons. Same conventions as Fig. 4j. **i.** Accuracy of an SVM classifier, similar to Fig. 2h, but trained on ABAB trials after removal of contingency neurons. The classifier no longer performs above chance on CDAB trials during the choice epoch.

Contingency neurons constituted 28.7% ± 1.7% of the choice-selective population (Fig. 5c**–d**). They tended to respond early in the choice epoch compared to the choice neuron population as a whole (Fig. 5e), suggesting that they may play a role in organizing or initiating the directed lick response. For example, the neuron in Fig. 5f was active during leftward licks, but only when preceded by context odors A¹^ˢᵗ^ or C¹^ˢᵗ^ (Cx1), corresponding to a specific contingency: A²ⁿᵈ → lick left. It is also noteworthy that these neurons are responsible for the convergence by context in the SVM classifier and PCA. Removal of these neurons from the PCA analysis abolished the divergence by context (Fig. 5h **and Supplementary** Fig. 4a); same for the context classification in the epoch following the test odor (Fig. 5i, *left*). The decoding accuracy improves linearly as a function of the number of contingency neurons included in the model (Fig. 5i, *right*). The contingency neurons were not required for choice decoding, as there are many choice (directed-lick) neurons (**Supplementary** Fig. 4b).

To determine whether this dual selectivity could arise by chance from independent context and choice tuning, we performed a permutation test. The observed number of contingency neurons significantly exceeded the null distribution derived from shuffled labels (**Supplementary** Fig. 4d). In contrast, we found only weak task-specific choice selectivity (e.g., neurons active in ABAB but not CDAB, likely reflecting training history rather than functional specialization; **Supplementary** Fig. 4e–g). Similarly, neurons that responded across mismatched context–stimulus pairings were rare and did not exceed chance levels (**Supplementary** Fig. 4h). Together these observations suggest that contingency neurons link contextual history to motor response, thus implementing the flexible SR-mapping.

## Discussion

Context-dependent decision-making is a hallmark of cognitive flexibility. It enables the brain to apply appropriate stimulus–response (SR) mappings based on internal goals or external conditions. At its core, this ability is compositional: the meaning of a stimulus depends on the context in which it occurs. Context guides a decision-maker to apply the appropriate strategy or behavior, contingent on desired outcomes. In binary choices, the same evidence may support opposite actions (e.g., approach vs. avoidance) ^19–21^ or modulate the vigor of a common action (e.g., foraging under abundance vs. scarcity) ^22–25^. In our task, the same test odor (A²ⁿᵈ) leads to different actions depending on whether it follows context odor C¹^ˢᵗ^ or D¹^ˢᵗ^. This study is a first step in our quest to understand how the anterior lateral motor cortex (ALM) implements such context-sensitive reconfigurations.

A previous study by Wu et al. examined this process using a delayed match-to-sample (DMS) task. Mice were presented with a sample odor (A¹^ˢᵗ^ or B¹^ˢᵗ^), followed by a test odor (A²ⁿᵈ or B²ⁿᵈ). Neurons in ALM exhibited selectivity for the sample odor, and optogenetic inactivation during the sample and delay periods impaired performance. Wu et al. proposed that the sample odor functions as a contextual cue that instructs ALM to apply a specific SR-mapping between the test odor and the directed lick response. This interpretation was supported by partial inactivation experiments, which suggested that ALM activity configures the mapping—not merely by retaining a memory trace of the sample odor for comparison, but by preparing the circuit to apply the appropriate transformation. The next step in this program is to understand the cellular and subcellular mechanisms by which context reinstates a learned SR-mapping. But before pursuing that goal, we aimed to rule out a simpler explanation: that ALM supports performance in the DMS task by enabling a sensory comparison between test and sample. To do so, we developed a task of identical structure that cannot be solved by same–different comparisons.

The CDAB task achieves this disambiguation directly, and the underlying ALM activity—revealed through calcium imaging—closely parallels the findings from the DMS task. The key difference is the emergence of “contingency neurons,” which encode specific combinations of context and action. We suspect these neurons were also present in the earlier study; they manifest clearly in ABAB trials interleaved in the 8T task. Whether such neurons arise in mice trained only on the ABAB task remains uncertain. But for our purposes, this is moot: the CDAB task provides the necessary leverage to investigate the mechanisms of context-dependent remapping.

We considered two possible strategies by which ALM might support this behavior. The first posits that ALM generalizes across context odors that instruct the same SR-mapping—for example, A¹^ˢᵗ^ and C¹^ˢᵗ^ might activate a shared neural representation of context Cx1. However, although we observed L2/3 neurons selective for each context odor, overlap between same-context odor pairs was no greater than between opposing contexts. A classifier trained to distinguish A¹^ˢᵗ^ from B¹^ˢᵗ^ could not distinguish C¹^ˢᵗ^ from D¹^ˢᵗ^, and vice versa. These results suggest that ALM maintains distinct representations for each context odor.

The second strategy proposes that ALM reconfigures local circuits on each trial, such that the same test odor drives different outputs depending on the context identity of the first odor. Our data support this view. Specifically, ALM circuits converge on shared motor outputs through a subset of “contingency neurons” that respond to specific combinations of context and test—for example, firing when A²ⁿᵈ follows C¹^ˢᵗ^ but not D¹^ˢᵗ^. These neurons are jointly selective for context (Cx1 or Cx2) and choice (left or right lick). They tend to be active early in the choice epoch, relative to the larger population of choice-selective neurons with similar preferences, and may help organize or initiate the resulting bout of licking. In contrast to neurons that separately encode context or choice, contingency neurons represent the SR-mapping itself.

An unresolved question is how context-dependent responses in contingency neurons are implemented at the circuit level. Two broad classes of mechanisms are possible, both of which assume that, by default, contingency neurons are unresponsive to excitatory input from test-odor–selective neurons. In one model, context delivers an instructive signal—perhaps targeted to dendritic compartments—that transiently enables the appropriate test-odor input to drive the neuron (Fig. 6a). In the other, the mapping is established through synaptic plasticity during learning, and context simply activates a preconfigured pathway (Fig. 6b). Both models imply a silent default state and raise important questions about how cortical circuits are dynamically configured to support flexible behavior. The instructive model, such as dendritic gating, offers a more flexible and efficient mechanism for remapping. Disentangling these alternatives will require further experimental and theoretical work. Understanding whether context establishes a transient gate or engages a fixed subcircuit will be essential to explain how cortical networks adapt their computations to task demands.

**Figure 6.**
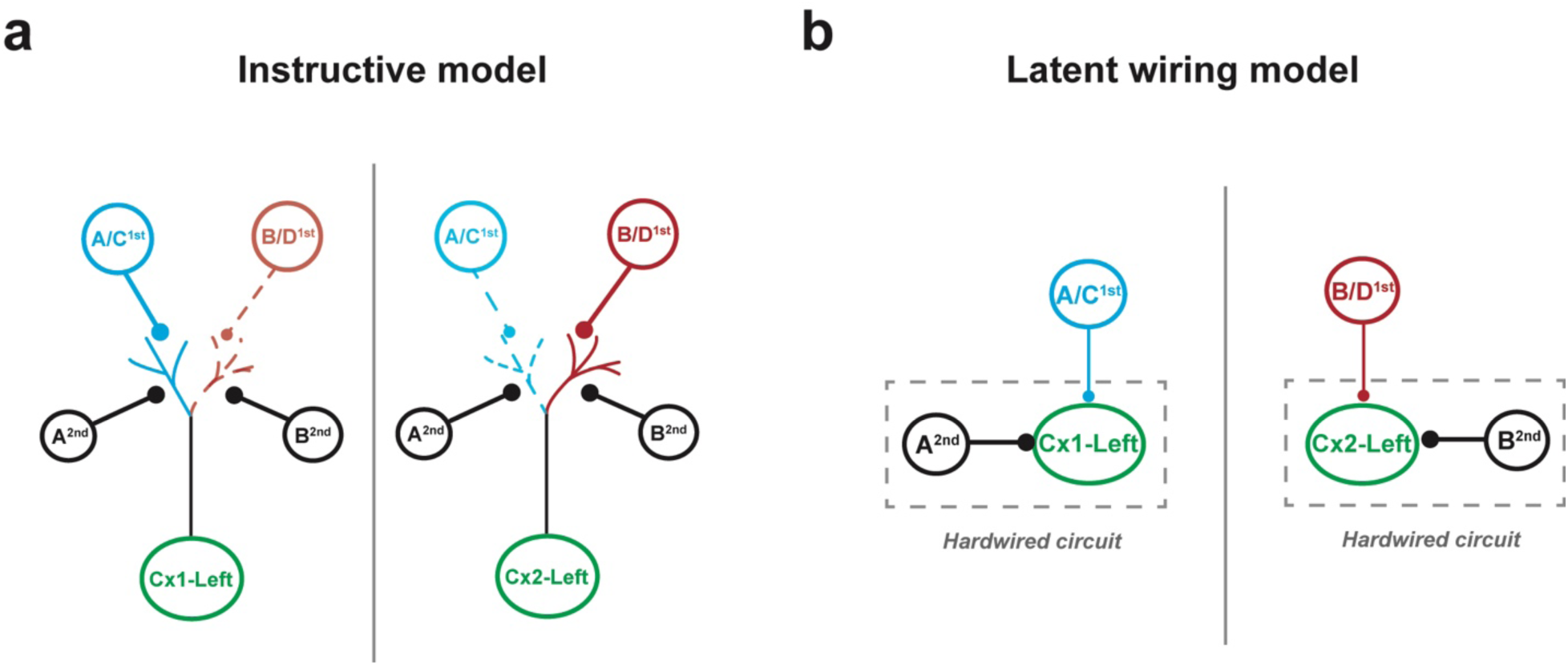
Two candidate mechanisms for context-dependent configuration of contingency neurons. For both mechanisms the contingency neuron is unreceptive by default to test odor input**. a**. Instructive model. *Left*, Cx1 left lick contingency neuron. Layer 2/3 A^1st^ and C^1st^ odor selective neurons prime the dendritic compartment receiving excitatory input from axons of A^2nd^ selective neurons. *Right*, Cx2 left lick neuron. Layer 2/3 B^1st^ and D^1st^ odor selective neurons prime the dendritic compartment receiving excitatory input from axons of B^2nd^ selective neurons. **B**. Latent wiring model. The connections are hard wired. *Left*, Cx1 left lick neuron (*latent*). The only test odor input is from neurons selective for A^2nd^. Superficial layer B^1st^ and D^1st^ odor selective neurons prime the neuron non-selectively, converting the neuron to a state of receptivity. *Right*, Cx2 left lick neuron (latent). The only test odor input is from neurons selective for B^2nd^. Superficial layer B^1st^ and D^1st^ odor selective neurons prime the neuron non-selectively; converting the neuron to a state of receptivity.

Our findings align with those of Kim et al.^10^, who found that preparatory activity in ALM varies with context, even when the planned movement is the same. This surprising result suggests that the neural substrate for a single action (e.g., a directed lick) can shift depending on task context. However, in their study, the two contingencies were learned sequentially, raising the possibility that distinct ALM populations reflected differences in training history rather than context per se. In contrast, our mice were trained to learn two contingencies in parallel (ABAB trials), followed by trials with new odor cues that preserved the same two mapping rules (CDAB trials). This design enabled us to disentangle the effects of sensory identity, learned mapping, and training sequence. The identification of contingency neurons—cells that respond selectively to a specific SR-mapping—provides direct evidence that context gates stimulus representations at the level of individual neurons, supporting flexible routing within a shared circuit. This may involve short-term synaptic modulation or dendritic gating^26,27^.

Several open questions remain. First, the source of contextual input to ALM is unknown. Although the orbitofrontal cortex (OFC) contains odor-selective neurons and projects to ALM, inactivation studies suggest it is not essential for task performance. The mediodorsal thalamus (MD), which is involved in motor planning and strongly connected to ALM, may be a more likely source. Second, the causal role of contingency neurons has not yet been tested. Targeted optogenetic perturbation could determine whether they are necessary for applying the correct SR-mapping.

These results also challenge standard interpretations of delayed match-to-sample (DMS) tasks. In the CDAB paradigm, context and test odors are drawn from non-overlapping sets, and the first odor is never repeated. As a result, mice cannot solve the task by comparing two sensory traces. This raises the possibility that other DMS paradigms—even those in primates^28–30^—may rely more on rule application than on sensory comparison^33^.

In summary, ALM supports flexible decision-making not by forming abstract representations of context, but by reconfiguring local circuitry to route the same sensory input to different motor outputs. This transformation, mediated by contingency neurons, allows the brain to construct behavior from composable elements. Such dynamic and modular operations may underlie core features of cognitive control.

## Resource availability

### Lead contact

Further information and requests for reagents may be directed to and will be fulfilled by the lead contact authors, Herbert Zheng Wu and Michael N. Shadlen.

## Materials availability

This study did not generate new, unique reagents.

No original code was generated in this study.

Any additional information required to reanalyze the data reported in this paper is available from the lead contact upon request.

## Acknowledgments

We thank Emma Ruimschotel-Quinn and Institute of Comparative Medicine (ICM) at Columbia University for their supportive work. We thank Lina Marcela Carmona, David C. Gruskin, and all members of the Shadlen lab for their constructive discussions throughout the study. We are grateful to Richard Axel for his inspiration and guidance throughout the project. The research was supported by the Howard Hughes Medical Institute (M.N.S.), the NIH Brain Initiative (M.N.S., R01NS113113), the Grossman Center for the Statistics of the Mind at Columbia University, the Air Force Office for Scientific Research (21RT0878), the NIMH (M.N.S., R01MH122513; H.Z.W., R01MH133039), and NIH/NIDA F32 (J.S., 5F32DA060751).

## Author contributions

J.S., H.Z.W., and M.N.S. designed the study. J.S. and M.N.S. wrote the original manuscript. J.S., H.Z.W., N.R., D.S.P., and M.N.S. edited the manuscript. H.Z.W. and J.S. performed animal surgery, image collection, and data analysis. P.S. and J.S. performed behavior tests and data analysis. D.S.P. and N.R. assisted with some experimental design and data analysis.

## Methods

### Ethics statement

Animal care and experiments were carried out in accordance with the NIH guidelines and approved by the Columbia University Institutional Animal Care and Use Committee (IACUC protocol # AC-AABI4554).

### Animals

This study is based on data from forty-three mice (both males and females, 2-8 months old). Twenty-six C57BL/6J (Jackson Laboratory, JAX 000664) mice were used for behavioral paradigm testing and training. Optogenetic inhibition experiments were performed using twelve VGAT-ChR2-EYFP mice (Jackson Laboratory, JAX 014548). Two-photon imaging data were collected from five Emx1-Cre+/;TITL-GCaMP6f+/;CaMK2a-tTA+ mice. These mice were created by crossing Ai93(TITL-GCaMP6f)-D;CaMK2a-tTA (JAX 024108) to Emx1-IRES-Cre+/+ (JAX 005628) line. Prior to training, animals were maintained on 12 h: 12 h light/dark cycle with food and water available *ad libitum*. Mice were water-restricted during the training and testing phases. Experimental sessions were 1-2 h, during which mice received 0.5-1.5 mL of water. Animals received supplemental water as necessary to maintain their body weights. Aseptic surgeries were carried out under ketamine (100 mg/kg)/ xylazine (10 mg/kg) or 1%–3% isoflurane anesthesia. Buprenorphine (0.05-0.1 mg/kg) and carprofen (5 mg/kg) were administered for postoperative analgesia.

### Behavior training

Before training, mice were implanted with a custom-made titanium head post^16^. The scalp and periosteum over the dorsal surface of the skull were removed, and a head post was placed on the skull, aligned with the lambda suture and cemented in place with C&B metabond (Parkwell, S380). After at least 2-3 days of recovery, animals were water-restricted and accustomed to head-fixation following procedures described in Guo et al.^16^ and then trained on a custom-assembled apparatus.

*CDAB task:* Odorants were delivered with a custom-made olfactometer and custom-written LabVIEW programs (National Instruments). (+)-a-pinene (odor A, Sigma-Aldrich, 268070), cis-3-hexen-1-ol (odor B, Sigma-Aldrich, W256307), (R)-(+)-limonene (odor C, Sigma-Aldrich, 183164), and methyl butyrate (odor D, Sigma-Aldrich, 246093) were chosen for their lack of innate valence to mice (Root et al., 2014) and their low adhesion to the surface of the olfactometer. The odorants were diluted 1:100 in mineral oil (Fisher Scientific, O121-1) and then loaded on syringe filters (GE healthcare, 6888-2527 or 6823-1327). The air flow was maintained at 1.0 L/min. We confirmed the rapid kinetics of these odorants with photo-ionization detector as previously described^6^. During the training phase, mice were presented with a context odor (1.0 s duration) and a test odor (1.0 s) separated by a delay epoch (1.0 s). After hearing an auditory ‘‘go’’ cue (0.1 s, 5 KHz pure tone), they were free to report their decision by licking to one of the two syringe ports positioned in front of their mouth during a 2.0 s response window. A water reward was delivered at the same port if the choice were correct. A choice is scored as correct only when the first two licks were on the correct port. If they first licked on the incorrect port, that trial was scored as an error. If they licked once on the correct port and then on the incorrect port, that trial was also scored as an ‘‘error.’’ Mice were punished by a brief timeout (3-8 s) when they made an error. If they did not lick during the ‘‘response window’’ or only licked once on the correct port, that trial was scored as ‘‘no choice.’’ Animals completed this training stage when they achieved a criterion of at least 80% correct for each trial type in a single session. After reaching this criterion, the context and test durations were reduced to 0.5 s, and the delay was increased to 1.5–2.5 s depending on the experiment (2.0 s in 2p Ca^2+^ imaging experiments, 1.5 or 2.5 s in optogenetic inactivation). Mice performed at 90% correct (interquartile range 88%–92%) when they entered the testing phase. The four odorants give rise to four unique pairs of context and test odors (CA, CB, DB, and DA), or trial types, and were randomly presented in each session. Two trial types (CA, DB) were rewarded on the left port and the other two (CB, DA) on the right.

*8T task:* Mice were first trained on the ABAB task, which included four trial types (AA, AB, BB, and BA), as described in Wu et al. Once they reached a criterion of at least 80% correct for each trial type, training transitioned to the CDAB task protocol as described above. After achieving the same performance criterion on the CDAB trials, mice were presented all eight trial types in a randomized and interleaved manner. Mice performed at 90% correct (interquartile range 88%– 92%) when they entered the 2p Ca^2+^ imaging phase.

### Optogenetic inhibition

The inhibition experiments were performed as previously described^6^. Briefly, animals were prepared with a clear skullcap to achieve optical access to ALM^16^. After removing the scalp and periosteum over the dorsal surface of the skull, a layer of cyanoacrylate adhesive (Krazy glue, Elmer’s Products Inc.) was directly applied to the intact skull. The entire skull was then covered with a thin layer of the clear dental acrylic (Lang Dental) with a head post cemented over the lambda suture. Before photostimulation sessions, the dental acrylic was polished (0321B, Shofu Dental Corporation) and covered with a thin layer of clear nail polish to reduce glare (Part No. 72180, Electron Microscopy Sciences). Light from a 473nm laser (MLL-FN-473-50mW, Ultralasers, Inc.) was directed to an optic fiber and split into two paths (FCMH2-FCL, Thorlabs, Ø200 mm Core, 0.39 NA). The two optic fibers were positioned over ALM on each hemisphere. The light transmission through the skullcap is ∼50% in average power, as measured directly with a light meter (PM100D, Thorlabs).

We used 40 Hz photostimulation with a sinusoidal temporal profile (3 mW average power as light reaches the skull, ∼1.5 mW on the brain surface). The photoinhibition inactivated a cortical area of ∼1mm radius. To reduce rebound excitation after laser offset, we included a 250 ms linear power ramp-down at the end of the photostimulation unless otherwise indicated. To prevent the mice from distinguishing photostimulation trials from control trials, a masking flash (40 Hz sinusoidal profile) was delivered with 470 nm LED (Luxeon Star) and LED driver (SLA-1200-2, Mightex Systems) in front of the animals’ eyes on all trials. The masking flash began at context odor onset and lasted until the end of test odor stimuli, covering the entire stimulus and delay epochs in which photostimulation could occur.

For the experiments in which we varied the duration of inactivation, photostimulation was either limited to (*i*) the entire sample plus delay epoch (Fig. 3b) or (*ii*) the 0.5 s sample epoch and the first 1.5 s of the 2.5 s delay epoch (D1, Fig. 3c). These inactivation trials were randomly interleaved to constitute 25% of all trials. Note that we did not repeat the control inhibition experiment where the context odor is not informative since the structure of the task will be identical to that in Wu et al. And we expect the performance was not affected by the ALM inhibition^6^.

### 2-photon Ca^2+^ imaging

Calcium imaging was performed on Emx1-cre+/;TITL-GCaMP6f+/;CaMK2a-tTA+ mice. A square craniotomy (2mm side) was made above left or right ALM, along the superior sagittal sinus and the inferior cerebral vein. The imaging window was constructed from three stacked layers of custom-cut coverglass (CS-3S, Warner Instruments) and cemented with C&B metabond. Animals were allowed 1-2 weeks of recovery before the imaging sessions began. Images were acquired with a Bruker Ultima 2P Plus two-photon microscope under bi-directional resonant galvo scanning mode, and collected green fluorescence detected with GAsP PMTs (Hamamatsu Model H10770PB-40) filtered by a Chroma ET52/40m filter. The light source was a femtosecond pulsed laser (Chameleon Vision II, Coherent). The objective was a 16X water immersion lens (Nikon, 0.8 NA, 3mm working distance). GCaMP6f was excited at 920nm and images (512 3 512 pixels, ∼820 mm x 820 mm field of view) were acquired at ∼30 Hz.

## QUANTIFICATION AND STATISTICAL ANALYSIS

### Ca^2+^ imaging data processing and analysis

The raw images were first motion corrected and regions of interest (ROIs) were detected and registered automatically with Suite2p^31^. For data analysis, we manually computed unfiltered DF/F traces as follows. We obtained the raw fluorescent trace of each ROI by applying the spatial component (ROI filter) on the image sequence. We then smoothed the raw trace in each trial with a 1 s averaging window and take the minimal fluorescence value in the inter-trial interval as the baseline. The DF/F signals were calculated by subtracting the baseline from the raw trace and dividing the difference by the baseline.

We classified the selective neurons as follows. We compared the means of the response to odors C and D during the context and delay epochs using a Wilcoxon signed-rank test. Each trial contributed a scalar value: the average DF/F signal from t = 0-2.5 s from onset of the context odor. Neurons were classified as context odor-selective if *p* < 0.01 (Fig. 2b, two-tailed, not corrected for multiple comparisons). Test odor (Fig. 2c) and choice (Fig. 2d) selectivity were determined similarly using time bins of 2.5-3.5 s and 3.0-5.0 s from context onset, respectively. Selectivity indices were computed from the same scalar values. To quantify stimulus and choice selectivity from Ca²⁺ dynamics, we computed ROC-based indices for each neuron using trial-averaged ΔF/F signals. For each comparison, we calculated mean fluorescence across a task-defined epoch and used the Wilcoxon rank-sum test to assess selectivity. The resulting U statistic was normalized by the product of trial counts to yield an ROC value equivalent to the area under the receiver operating characteristic curve (AUC). Final ROC values were scaled to range from –1 to 1, with positive values indicating preference for one condition and negative values for the other. Neurons were classified as contingency-specific when 1) they are choice-selective and 2) their Ca^2+^ response was significantly different across two contexts within the preferred choice.

The standardized odor and choice selectivity (Fig. 2e) were computed by dividing the absolute value of the difference between the mean DF/F responses to odor C and D (or lick-left and lick-right) by the common standard deviation: the standard deviation is the square root of the sum of the variances of the DF/F responses to odors C and D (or lick-left and lick-right).

To determine the duration of the calcium response (Fig. 2f), we first computed the mean and standard deviation of the baseline (the epoch before context odor onset). Calcium transients were then identified as any response greater than 2 standard deviations away from the baseline and lasting for at least 100 ms. The peak time is the time of the maximum calcium response. Response duration is the median duration of each neuron’s response. Scatter is defined as the interquartile range of the peak response times across trials.

For the decoding analysis (Fig. 2h, 4h, 5i, and **Supplementary** Fig. 3f and 5a), we trained a support vector machine (SVM)^32^ using the Ca^2+^ responses to classify context odor, test odor, or choice. The calcium responses were computed as the average DF/F in a sliding bin of 100 ms at 100 ms steps. In Fig. 4h and 5i, we trained the model using 80% of the ABAB trials and cross validated on the rest 20% of the trials. The same model was used to decode the variables in all the CDAB trials. The training and testing data were reversed in **Supplementary** Fig. 3f. To assess the temporal stability of neural representations extracted by the decoder, we computed the cosine similarity between linear SVM weight vectors trained at different time bins. Cosine similarity was defined as the dot product of two normalized weight vectors and reflects the angular similarity between feature representations, independent of their magnitude. We applied a sliding window (5 time bins wide) to smooth the weight vectors before calculating pairwise similarities. To quantify whether representations were more stable within an epoch (e.g., before test odor onset) compared to across epochs (e.g., before vs. after), we averaged cosine similarities within each region of the similarity matrix (before-before, after-after, and before-after). We then performed a Monte Carlo permutation test to determine whether the observed differences in mean similarity (e.g., before-before vs. before-after, and after-after vs. before-after) were statistically significant. For each comparison, we generated a null distribution by randomly shuffling group labels (1,000 permutations) and recalculating the mean difference for each shuffle. Two-tailed p-values were computed as the proportion of permuted differences that exceeded the observed difference in absolute value.

For PCA analysis, DF/F values between the context odor onset and delay offset were used in **Supplementary** Fig. 3g and that between test odor onset and choice window offset were used in Fig. 4j and Fig. 5h. The three principal components that explained the most variance were used to visualize the neural trajectory.

### Statistical

Statistical analysis was performed using MATLAB. Sample sizes were chosen based on our previous work with similar techniques^6^. No outlier data were identified or removed. Data sets with normal distributions were analyzed for significance using paired or unpaired Student’s two-tailed t test or analysis of variance (ANOVA) measures followed by Tukey’s post hoc test. Data sets with non-normal distributions were analyzed using Wilcoxon signed-ranked test or Kruskal–Wallis test. Correlations were examined using Pearson’s r test. Data are expressed as mean ± s.e.m throughout. n indicate number of mice unless indicated otherwise. The significance threshold was set at *p* < 0.05.

---

**Supplementary Figure 1.**
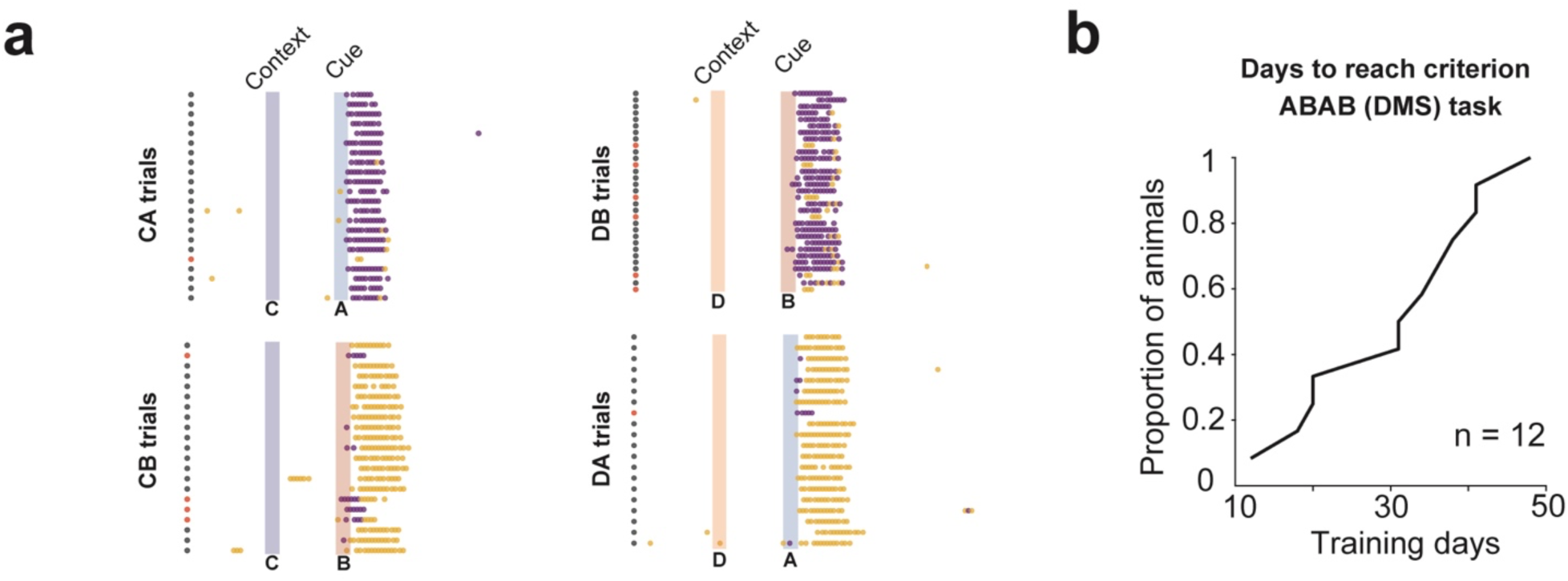
a. Days required for behavioral training for the ABAB task paradigm (n = 12, 30.8 ± 3.2 days). **b**. Example session with licking timestamps (black: correct trials, red: error trials; purple: licking left, yellow, licking right).

**Supplementary Figure 2.**
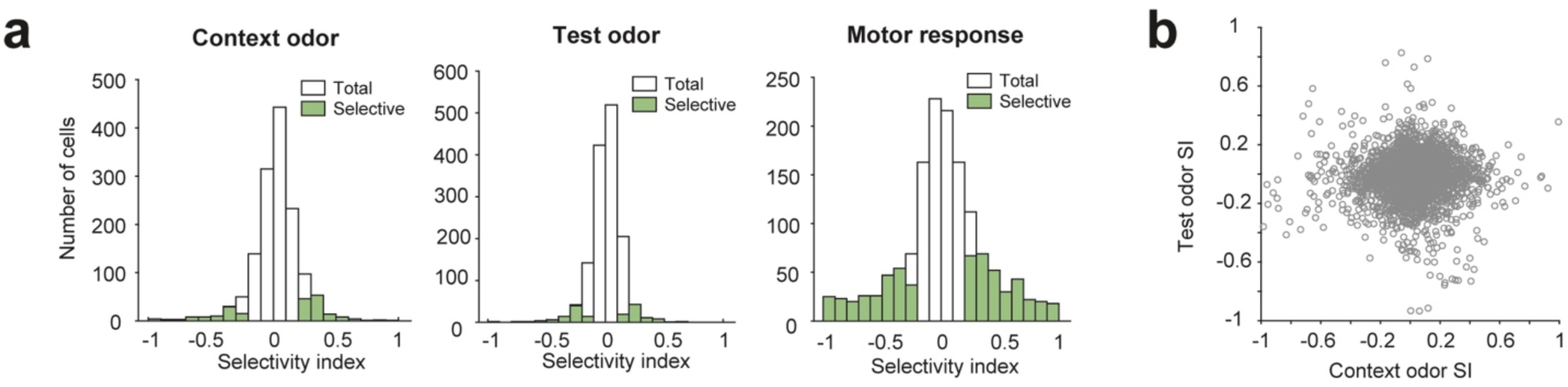
a. Summary of the selectivity index in each cell type in the CDAB task. b. Selectivity indices of ALM neurons for context and test odor. Pearson’s correlation R = 0.03 (P= 0.1).

**Supplementary Figure 3.**
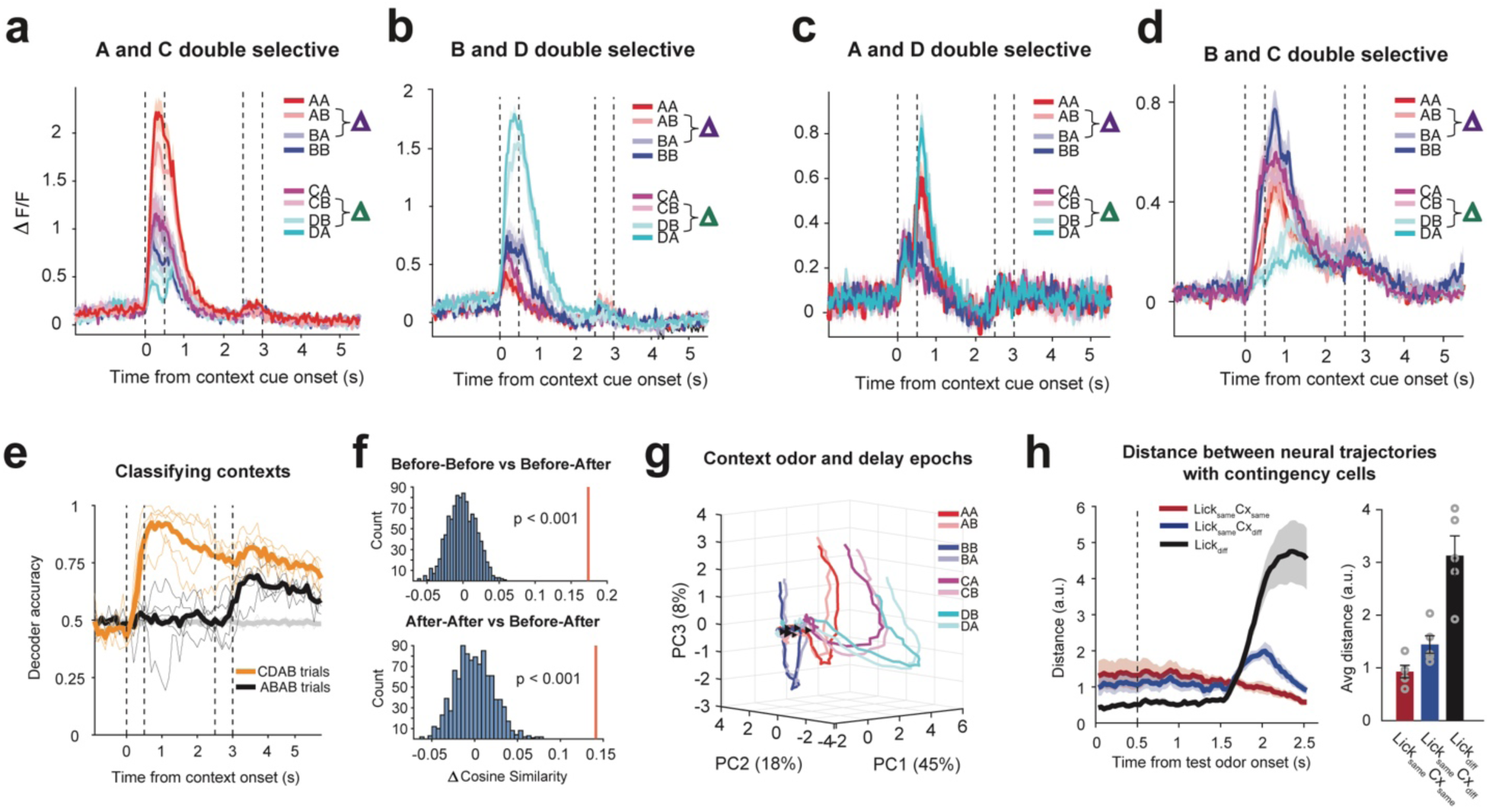
a-d. Example double selective neurons with Ca2+ responses in all trial types corresponding to the three neurons in Fig. 4d-g. e. SVM models trained using neural activity in the CDAB trials and decode contexts in the ABAB ones (black). Shuffled data is shown in grey and cross validated decoder accuracy is shown in orange. f. Monte Carlo permutation test to assess the cosine similarity between before and after the test odor onset. The difference between before-after is statistically larger than those within the same time block (before-before or after-after). g. Neural trajectory in a 3D state space showing the divergence of context odors (triangles indicate the test odor offset). **h.** Euclidean distance between the PCA neural trajectories in three groups: trials in the same lick direction and context (red); trials in the same lick direction but different context (blue); and trials in different lick directions (black).

**Supplementary Figure 4.**
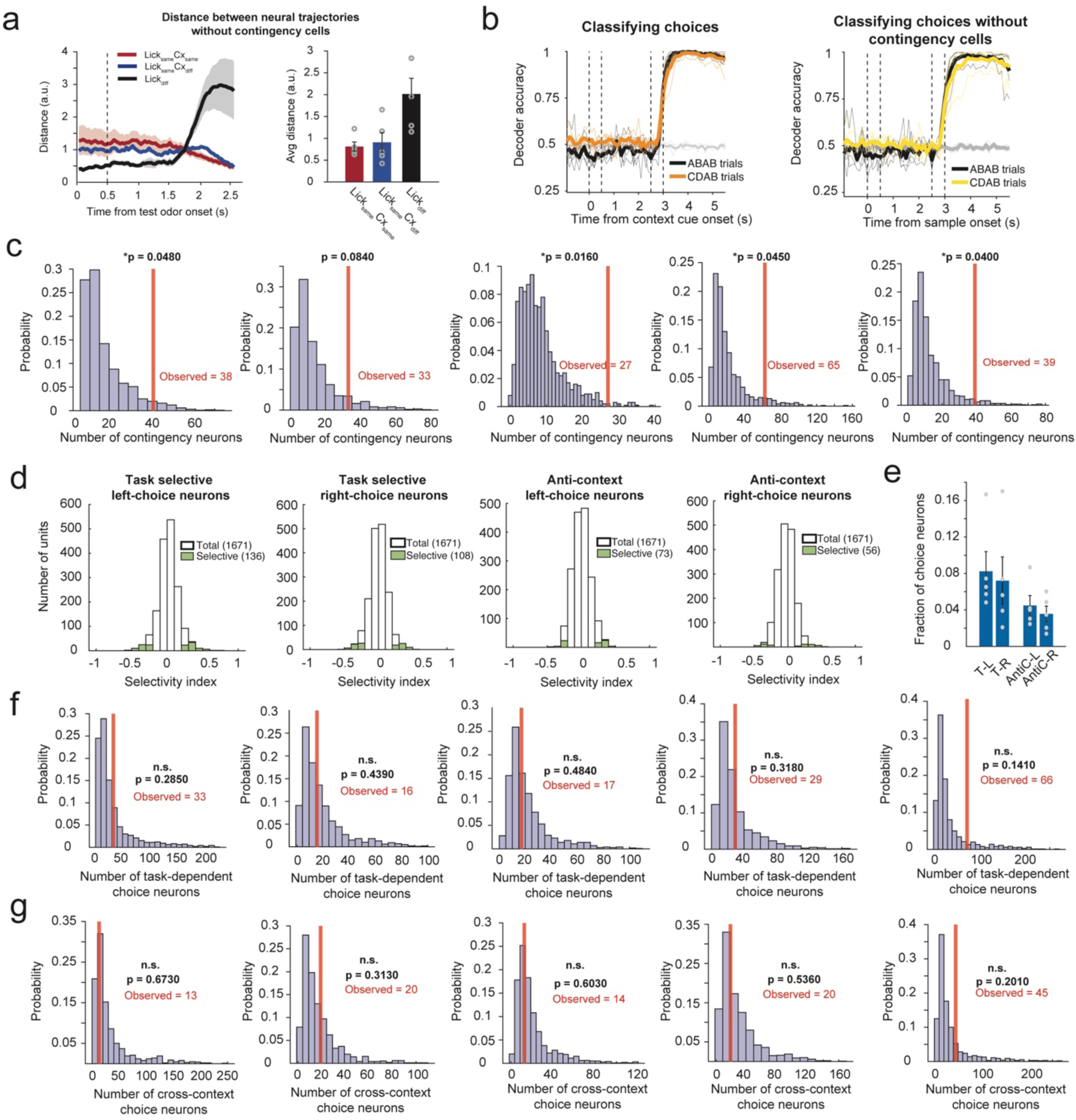
a. Euclidean distance between the PCA neural trajectories without contingency cells in three groups: trials in the same lick direction and context (red); trials in the same lick direction but different context (blue); and trials in different lick directions (black). b. SVM models trained using neural activity in the ABAB trial and decode choice with (left) or without (right) contingency cells. c. The amount of contingency cells are above the chance level (*p < 0.05, n = 1000 iterations, permutation test). Each plot is one individual animal (histogram showing the distribution of 1000 iterations of simulated overlap of contingency neurons; red line is the number of contingency in our data). d. The SI distribution of task-dependent and anti-context choice neurons. e. Percentage of each type of choice neurons. Permutation test to access the amount of task-dependent choice neurons (f) and cross-context choice neurons (g) are above the chance level (n.s., p > 0.05, n = 1000 iterations).

## Footnotes

1 We use X^1st^ as a place holder for either context odor

